# Anatomy-guided Inverse-phase-encoding Registration Method for Correcting Susceptibility Artifacts in Sub-millimeter fMRI

**DOI:** 10.1101/779272

**Authors:** Soan T. M. Duong, Son L. Phung, Abdesselam Bouzerdoum, Harriet G. Boyd Taylor, Alexander M. Puckett, Mark M. Schira

**Affiliations:** School of Electrical, Computer and Telecommunications Engineering, University of Wollongong; School of Psychology, University of Wollongong, Australia; College of Science and Engineering, Hamad Bin Khalifa University, Qatar; School of Psychology, University of Queensland, Australia; Queensland Brain Institute, University of Queensland, Australia

**Keywords:** Susceptibility artifact, echo planar imaging, high-resolution fMRI, inverse phase-encoding, anatomy-guided

## Abstract

Echo planar imaging (EPI) is a fast and non-invasive magnetic resonance imaging (MRI) technique that supports data acquisition at spatial and temporal resolutions suitable for brain function studies. However, susceptibility artifacts are unavoidable distortions in EPI. These distortions are especially strong in high spatial resolution images and can lead to misrepresentation of brain function in fMRI experiments. A common method for correcting susceptibility artifacts is based on a registration scheme which uses two EPI images acquired using identical sequences but with inverse phase-encoding (PE) directions. In this paper, we present a new method for correcting susceptibility artifacts by integrating a T1-weighted (T_1*w*_) image into the inverse-PE based registration, since the T_1*w*_ structural image is considered as a ground-truth measurement of the brain. Furthermore, the T_1*w*_ image is used as a criterion to select automatically the regularization parameters of the proposed image registration. Evaluations on two high-resolution EPI-fMRI datasets, acquired at 3T and 7T scanners, confirm that the proposed method provides more robust and sharper corrections and runs faster compared with two other state-of-the-art inverse-PE based susceptibility artifact correction methods, *i.e.* HySCO and TOPUP.

## 1. Introduction

Functional Magnetic Resonance Imaging (fMRI) indirectly estimates the changes in cortical activity, most often by measuring the Blood Oxygenation-Level Dependent (BOLD) signal (Ogawa et al., 1990). fMRI allows researchers and medical practitioners to non-invasively examine not only the structure but also the function of the human brain, and hence, fMRI has become widely used in clinical and research settings. At present, fMRI images are almost exclusively acquired using the EPI technique because of its fast temporal imaging capability. For example, EPI takes 1 to 3 seconds to scan a volume compared to about 5 minutes for most other MRI techniques. This capability enables EPI to record rapid changes in brain activity.

Despite its speed, EPI is prone to distortions due to local field inhomogeneities, which are caused by the difference in magnetic susceptibility of various imaged tissues (e.g., fat versus blood) (Ludeke et al., 1985; McRobbie et al., 2003). The field inhomogeneities affect the spatial encoding of the signal. Consequently, they degrade the acquired images by geometrical deformations (stretching and compressing) and intensity modulations (Chang and Fitzpatrick, 1992). These distortions are known as susceptibility artifacts (SAs). The SAs are more severe at high field strengths (Ogawa et al., 1990; Polimeni et al., 2018) and in rapid imaging techniques such as EPI (Schmitt, 2015; Ludeke et al., 1985). These artifacts can be easily seen in the interface regions, particularly between the cerebral cortex and non-brain areas (McRobbie et al., 2003). In practice, SAs are most noticeable along the PE direction. Pertinently, they appear reversed in two EPI images acquired using identical sequences but with inverse PE directions^1^ (Jezzard and Balaban, 1995; Hutton et al., 2002; Holland et al., 2010).

The SAs disrupt the geometric correspondence between functional and anatomical data. This disruption subsequently leads to misplacements of detected activation patterns in fMRI studies. Currently, correcting SAs in fMRI is often avoided for two main reasons. First, fMRI data have a spatial resolution of 1mm^3^ or greater, where SAs are generally not severe enough to cause a significant problem. However, the impact of the SAs is much more significant in high spatial resolution (sub-millimeter) fMRI, which has become widely used. Second, existing SA correction methods tend to blur the corrected images (Polimeni et al., 2018), which contradicts the goal of acquiring a higher spatial image resolution.

The work presented in this paper aims to correct SAs in EPI-fMRI images, especially those with sub-millimeter resolution. We propose to integrate a T_1*w*_ structural image into a state-of-the-art susceptibility artifact correction (SAC) scheme, known as hyper-elastic susceptibility artifact correction (HySCO) (Ruthotto et al., 2012). The justification is that the T_1*w*_ image captures relatively well the actual shape and size of the tissue. Thus, it is widely considered as a *gold standard* representation of a subject’s brain anatomy. It is routinely acquired for every subject participating in fMRI studies because of the high level of tissue contrast between white and gray matter (Polimeni et al., 2018), and hence is readily available. We call the proposed method Anatomy-guided Inverse-phase-encoding Susceptibility Artifact Correction, or AISAC.

The research contributions of this paper can be highlighted as follows. First, a new T_1*w*_-based regularization term is introduced to the HySCO objective function to improve the quality of the corrected image with respect to the brain structure captured by the T_1*w*_ image. Second, the regularization parameters of the registration problem are selected automatically through a Bayesian optimization framework with a Gaussian process prior. This approach is because choosing the best regularization parameters is a critical step in solving the SAC optimization problem. Furthermore, we evaluate the performance of the proposed method and compare it with existing SAC methods using two high-resolution EPI-fMRI datasets: one with an isotropic resolution of 1 × 1 × 1 mm^3^ acquired by a 3 Tesla (T) scanner, and the other with a resolution of 0.833 × 0.833 × 0.810 mm^3^ acquired by a 7T scanner.

The remainder of this paper is organized as follows. Section 2 presents the related work and the general mathematical framework of the inverse-PE based correction method. Section 3 introduces our proposed method. Section 4 presents experiments and analysis of the proposed method and the related methods. Finally, Section 5 summarizes our work.

## 2. Related work

In this section, an overview of the existing SAC methods is presented in Subsection 2.1. The inverse-PE SAC formation is then described in Subsection 2.2. Finally, the HySCO method is discussed in Subsection 2.3.

### 2.1. Susceptibility artifact correction methods

Several SAC methods have been proposed for multiple types of MRI, such as structural MRI, diffusion-weighted MRI (DWI), and fMRI. In general, they can be divided into four categories: (i) fieldmap based; (ii) point spread function (PSF) based; (iii) image registration based; and (iv) inverse phase-encoding (PE) based methods. Table 1 summarizes the SAC methods discussed below.

**Table 1:** Representative methods for correcting SAs (Seq. mod. = pulse sequence modified).

| Category | Seq. mod. | Authors | Year | Datatype | Description |
| --- | --- | --- | --- | --- | --- |
| Fieldmap | No | Jezzard and Balaban | 1995 | EPI | Use the fieldmap derived from complex images acquired by different TEs to unwarp the distorted images. |
|  |  | Reber et al. | 1998 | EPI | Smooth the displacement derived by the method in Jezzard and Balaban (1995) using a 2D Gaussian kernel to increase the signal-to-noise ratio. |
|  |  | Hutton et al. | 2002 | fMRI | Derive the fieldmap from EPI images acquired with different TEs. |
|  | Yes | Kadah and Hu | 1997 | EPI | Use the fieldmap to rewind the additional accumulated phase in k-space (called SPHERE). |
|  |  | Wan et al. | 1997 | EPI | Calculate the fieldmap using a set of reference scans generated by turning off the PE gradient of the EPI pulse sequence. |
|  |  | Chen and Wyrwicz | 1999 | EPI | Incorporate a set of fieldmaps by the multi-channel modulation algorithm to obtain corrected images. |
|  |  | Techavipoo et al. | 2008 | EPI | Derive the fieldmap from EPI images with modified k-space trajectories. |
| Point spread function | Yes | Robson et al. | 1997 | EPI | Measure the PSF by an EPI sequence with added PE gradients, constant time but variable magnitude. |
|  |  | Munger et al. | 2000 | EPI | Unwarp the distorted image given the measured PSF by a conjugate gradient algorithm. |
|  |  | Zeng and Constable | 2002 | EPI | Correct both the intensity and geometric distortions in EPI images by measured PSF as in Robson et al. (1997). |
|  |  | Zaitsev et al. | 2004 | EPI | Measure the PSF by integrating a parallel imaging technique into the acquisition. |
| Image registration | No | Kybic et al. | 2000 | EPI | Register distorted EPI images by modelling the displacement with splines and using the SSD similarity measure. |
|  |  | Studholme et al. | 2000 | fMRI | Register EPI images using a multimodality non-rigid registration algorithm with log-intensity measure. |
|  |  | Wu et al. | 2006 | fMRI | Register distorted images based on Thirion's demons. |
| | | Wu et al. | 2008 | EPI | Register distorted EPI images to a $T_{2w}$ using mutual information. |
| Inverse-PE | No | Chang and Fitzpatrick | 1992 | Structural MRIs | Introduce the theoretical justification of the correction using inverse phase-encoded images; correct each 1D image along the PE direction independently by finding pairs of corresponding points in the given two images. |
|  |  | Andersson et al. | 2003 | DWI | Model the displacement as a function of discrete cosine basis functions (called TOPUP). |
|  |  | Holland et al. | 2010 | fMRI | Model the inverse-PE SAC as a diffusion registration problem. |
|  |  | Ruthotto et al. | 2012 | DWI | Introduce an additional non-linear regularizer into the diffusion regularized problem (called HySCO). |
| | | Irfanoglu et al. | 2015 | DWI | Incorporate a $T_{2w}$ image into the inverse-PE registration. |

*Fieldmap based* SAC methods estimate phase dispersions caused by the field inhomogeneity. The estimated phase dispersion over the entire scanned view is called the fieldmap. The corrected images can be obtained by unwarping distorted images (Jezzard and Balaban, 1995; Reber et al., 1998), or rewinding the additional accumulated phase in k-space (Kadah and Hu, 1997), thereby obtaining the corrected image. There have been multiple approaches to estimate the fieldmap. The original (and simple) approach derives the fieldmap from two complex MRI images acquired by different values of echo time (TE) (Hutton et al., 2002). The advanced approach, which can produce the fieldmap quickly, requires modified MRI sequences (Wan et al., 1997; Chen and Wyrwicz, 1999; Techavipoo et al., 2008). The main limitation of unwarping in the image space is the lack of intensity correction. Rewinding in k-space allows both geometric and intensity corrections but typically requires customized sequences.

*Point spread function based* SAC methods consider an acquired image as a convolution between the “true” image with a PSF. By estimating the PSF of the system, the undistorted image can be reconstructed. A PSF estimation technique based on constant time imaging was first introduced by Robson et al. (1997) for correcting EPI distortions and quantifying the MRI degradation. Subsequently, the PSF estimation was adopted to correct EPI distortions by Munger et al. (2000); Zeng and Constable (2002). A further optimized PSF estimation was proposed by integrating parallel imaging into the acquisition to correct distortions faster and more reliably, even at high field strengths (Zaitsev et al., 2004). PSF-based SAC methods can correct both geometric distortions and intensity modulations, however, they require the MRI scanner to support configurable MRI sequences.

*Image registration based* SAC methods map the distorted EPI images to a reference image using a non-rigid model. These methods usually estimate displacements in the image volume so that the unwarped image is morphologically matched to the reference image. These methods have several variants, based on the similarity measure between the EPI and reference images, *i.e.* the sum of squared differences (SSD) (Kybic et al., 2000), log-intensity metric (Studholme et al., 2000), and mutual information (Wu et al., 2006, 2008). An advantage of this approach is that it they do not require additional scans as the fieldmap-based methods do. However, methods in this class typically lack intensity distortion corrections and depend strongly on the constraints and parameters of the registration algorithms.

*Inverse phase-encoding based* SAC methods utilize two inverse-PE images to estimate the displacement field over the image domain. The corrected images are obtained by unwarping the distorted images by the estimated displacement field. Chang and Fitzpatrick (1992) initially introduced the theoretical justification of correcting the SAs using inverse-PE images. They then proposed a “cumulative line-integral” method to find the corresponding points, which are used to determine the displacement in two corresponding lines along the PE direction of the given inverse-PE images. Bowtell et al. (1994) implemented the original inverse-PE method for 2D EPI. The corrections of the method proposed by Chang and Fitzpatrick (1992) are not smooth since the method estimates the displacement in each line along the PE direction independently, without considering surrounding lines. To estimate the displacement field, Andersson et al. (2003) proposed an alternative approach by considering the displacement at a pixel as a function of discrete cosine basis functions to construct an objective function; this method is called TOPUP and is integrated into the FSL package^2^. Holland et al. (2010) integrated the inverse-PE approach into a registration framework to correct SAs. Ruthotto et al. (2012, 2013) combined the registration framework and a constraint inspired by the hyper-elastic image registration to achieve more realistic corrections; this method is called HySCO, and its implementation is included in the SPM12 toolbox^3^. Another approach combines an independent image, specifically a T_2*ω*_ image, into the inverse-PE registration to regularize corrections (Irfanoglu et al., 2015). Inverse-PE based SAC methods can correct both geometric and intensity distortion. They outperform fieldmap and image registration based methods in terms of geometrical correction fidelity, as shown in Hong et al. (2015). The inverse-PE based approach is the most common SAC method, for example being used to correct the fMRI data in the biggest MRI neuroimaging dataset - the Human Connectome Project (HCP) (Essen et al., 2012). However, registering corrected images from two inverse-PE images introduces more constraints than typical registration algorithms, for example the smoothness of the displacement field and the alignment of the correction to the structural image. The inverse-PE methods may produce less meaningful and blurred corrections due to the lack of suitable constraints.

Existing methods have been designed mostly for DWI images but rarely for fMRI. These methods either require a long scanning time or correct only spatial distortions. Furthermore, they are often inadequate at correcting SAs in high-resolution fMRI, where the distortions are more severe than in low-resolution images.

### 2.2. Distortion model in the presence of the field inhomogeneity

Let *E* be the 3D ideal image, and *I* be an acquired (distorted) image. As shown in Chang and Fitzpatrick (1992); Studholme et al. (2000); Holland et al. (2010), the distortion model in the presence of field inhomogeneity *B* in the image domain is

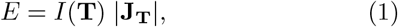

where **T** is the non-rigid transformation operator of coordinates from image *E* to image *I*, and **J_T_** is the Jacobian matrix of the transformation **T**. As shown in Holland et al. (2010); Ruthotto et al. (2012), the transformation **T** at any 3D point **p** in *E* can be written as **T**: **p** ↦ **p** + *B*(**p**)**v**, where **v** denotes the known distortion direction (*i.e.* the PE direction). In practice, the applied PE gradient is considered to be along the first dimension, hence **v** = (1, 0, 0). Let *∂*_**v**_(*B*(**p**) denote the directional derivative of field *B* at point **p** along the direction **v**. The Jacobian matrix of the transformation **T** at point **p** is

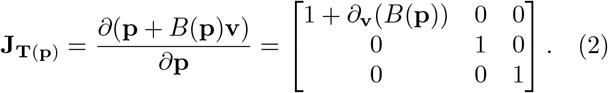

The distortion model in Eq. (1) can be rewritten as

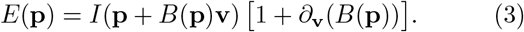

Here, the term 1 + *∂*_**v**_(*B*(**p**) denotes the intensity modulation. The term **p** + *B*(**p**)**v** denotes the geometric displacement of the acquired image. In other words, point **p** in ideal image *E* is shifted to point **p** + *B*(**p**)**v** in acquired image *I*. Since *B* causes the voxel shifting in the acquired image, *B* is called the *displacement field*, and **p** + *B*(**p**)**v** is known as the deformation at point **p**. Fig. 1 illustrates the distortions caused by the displacement field. The ideal image in Fig. 1(a) under the displacement field in Fig. 1(b) is distorted, as shown in Fig. 1(c). It is worth noting that we work with 3D images; however, for simplicity, 2D images are presented throughout this paper.

**Figure 1:**
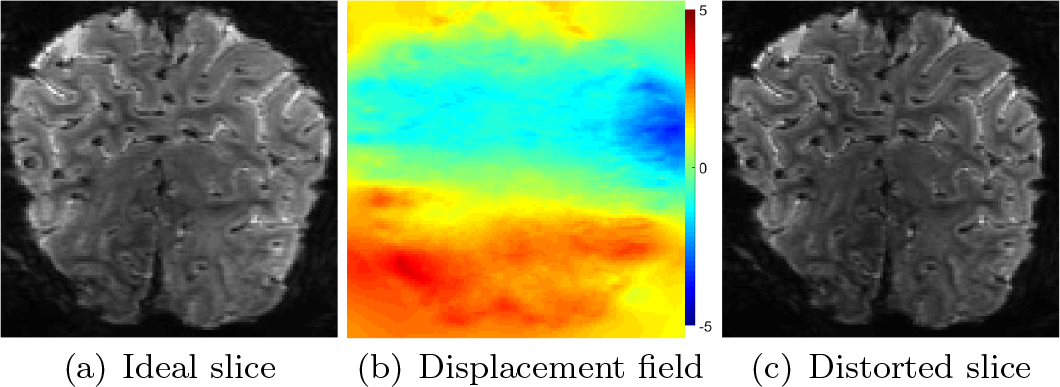
A 2D illustration representing the susceptibility-induced physical distortions. The displacement field is along the PE (horizontal) direction and is expressed in terms of the number of voxels shifted.

For two images *I*_1_ and *I*_2_ of the same subject’s brain region acquired using an identical sequence but with opposite blips, field inhomogeneities while acquiring these images have flipped signs. Let *B* be the field inhomogeneity, and **v** be the PE direction in acquired image *I*_1_. The field inhomogeneity and the PE direction in image *I*_2_ are *B* and −**v**, respectively. By applying the model in Eq. (3), the corrected images *E*_1_ and *E*_2_ can be described as

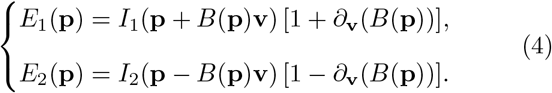

For notational simplicity, hereinafter *X*_**p**_ will refer to the intensity of image *X* at location **p**.

### 2.3. Hyper-elastic susceptibility artifact correction

Recall that the inverse-PE approach estimates the displacement field *B* based on two images *I*_1_ and *I*_2_ acquired using an identical sequence but with opposite blips. Field *B* is estimated such that two corrected images *E*_1_ and *E*_2_ are as similar as possible. The estimated B is then used to unwarp the distorted images *I*_1_ and *I*_2_ based on Eq. (4).

The hyper-elastic susceptibility artifact correction method proposed by Ruthotto *et al.* (Ruthotto et al., 2012) uses the inverse-PE approach to correct SAs. To estimate *B*, Ruthotto *et al.* minimized the SSD-based dissimilarity between unwarped images *E*_1_ and *E*_2_ (Holland et al., 2010; Ruthotto et al., 2012):

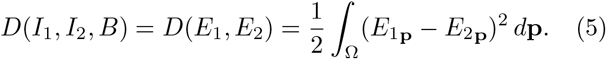

Finding *B* by minimizing the distance function *D*(*I*_1_, *I*_2_, *B*) is categorized as an ill-posed problem (Holland et al., 2010; Ruthotto et al., 2012). Thus, prior knowledge about the smoothness of the displacement field and invertibility of the geometrical transformation was introduced to regularize *B* (Ruthotto et al., 2012). To enforce the smoothness of the displacement field, a Tikhonov 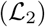 regularizer *S*^diff^ was integrated into the objective function (Holland et al., 2010):

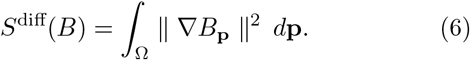

To satisfy the invertibility of the transformation, the Jacobian matrix of the geometric transformation in Eq. (4) must be invertible. In other words, Jacobian determinants must be positive for all **p** ∈ Ω. Chang and Fitzpatrick (1992) demonstrated that this constraint could be expressed as −1 ≤ *∂*_**v**_(*B*_**p**_) ≤ 1, for all **p** ∈ Ω.

Ruthotto et al. (2012), inspired by the control of volumetric change in hyper-elasticity (Burger et al., 2013), introduced an additional non-linear term *S*^hyper^ to the objective function:

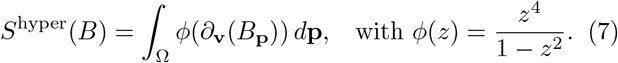

Collectively, Ruthotto et al. (2012) proposed the objective function:

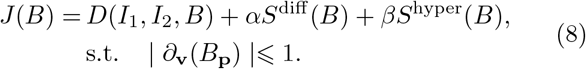

The positive and user-defined regularization parameters *α* and *β* represent the trade-off between the smoothness and the elasticity of the displacement field *B*.

The HySCO method estimates *B* by minimizing the objective function *J*(*B*) in Eq. (8), then generates the corrected images using Eq. (4). HySCO can provide corrected images with high similarity; however, the result is blurry and may not be reasonable in terms of the brain structure. For example, Fig. 2(a) shows the estimated deformation grid^4^ by HySCO and Fig. 2(b) shows the corrected image. The corrected image often contains blur trails (see areas denoted by red arrows). These artifacts are likely due to the over-deformation in estimated field *B* since there is no constraint regarding brain anatomy.

**Figure 2:**
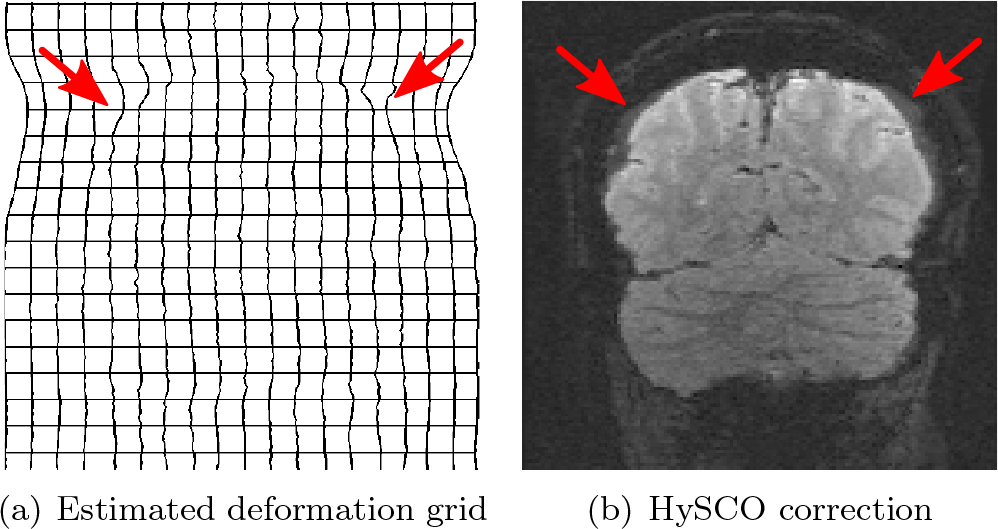
An example of HySCO results with additional artifacts (red arrows) on the top of the brain.

## 3. Anatomy-guided inverse-PE SAC

The T_1*w*_ structural image, which has different contrast properties (modality) to EPI-fMRI images, is considered as the ground-truth anatomy of the brain. Here, we utilize the brain anatomy provided by the T_1*w*_ image to introduce a robust and accurate SAC method for high-resolution EPI-fMRI images. More precisely, the T_1*w*_ image is used for two purposes. The first is introducing a new anatomy-guided regularization term for the registration problem. The second is selecting the regularization parameters automatically. The two aspects of our proposed method are in turn described in the following subsections.

### 3.1. AISAC registration

The inverse-PE correction problem integrated with a T_1*w*_ structural image can be formulated as finding the displacement field *B* such that the corrected (unwarped) images *E*_1_ and *E*_2_ satisfy two criteria: (i) be as similar to each other as possible and (ii) be the best fit with the brain anatomy provided by the T_1*w*_ image. Ruthotto et al. (2012) proposed the objective function in Eq. (8), which satisfies the first criterion. We introduce an anatomy-guided regularization term, which satisfies the second criterion, and add it into the objective function. More precisely, the regularization term measures the dissimilarity between the multimodal images, *i.e.* T_1*w*_ and EPI-fMRI. Thus, minimizing the proposed objective function is equal to minimizing the dissimilarity between the corrected EPI-fMRI images and between corrected images and the T_1*w*_ image. To get the registration method as accessible as possible, we describe the mathematical framework in the rest of this subsection.

The proposed regularization term is based on the normalized gradient field, which has been proven to be well-suited for the multi-modal registration problem (Haber and Modersitzki, 2007). The NGF measure, at any point in an image, reveals the intensity change and its direction. Let ∇*X*_**p**_ be the gradient at point **p** of image *X*, and *ɛ* be a user-defined parameter. As shown in Haber and Modersitzki (2007), the NGF measure at point **p** is defined as

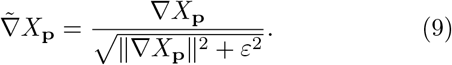

The difference between two images *X* and *Y* can be measured using the angles formed by NGF vectors at all points in the image domain. Accordingly, the NGF-based distance between two images *X* and *Y* is defined as

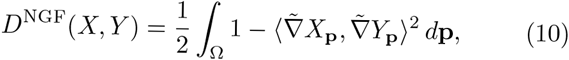

where ⟨·,·⟩ denotes the dot-product operator. The value of *D*^NGF^(*X, Y*) is positive. The smaller the value of *D*^NGF^(*X, Y*) is, the more similar are the two images.

Let *A* denote the T_1*w*_ image. We introduce the anatomy-guided regularization term as the sum of the NGF-based distances of image *A* with each unwarped image of *I*_1_ and *I*_2_ under the displacement field *B*

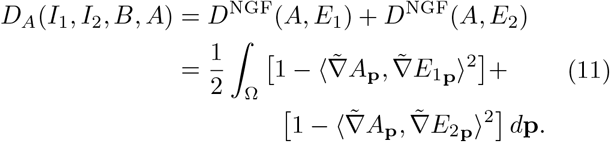

To summarize, we introduce a new objective function:

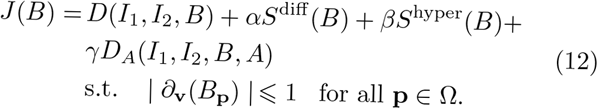

The displacement field is found by minimizing *J*(*B*) in Eq. (12). The positive and user-defined regularization parameters *α*, *β*, and *γ* represent the trade-off between the similarity of the corrected images, the smoothness of *B*, the elasticity of the displacement, and the similarity to the T_1*w*_ image of corrected images.

Here, the Gauss-Newton method is used for minimization. This method starts with an initial guess of *B*, e.g. *B*^(0)^ ≡ **0**. The next estimate of *B* is computed iteratively as

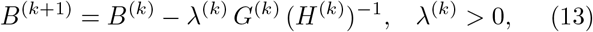

where superscript *k* is the iteration number, λ^(*k*)^ is the learning rate, and *G*^(*k*)^ and *H*^(*k*)^ are the approximate gradient and Hessian of the objective function *J*, respectively.

A small learning rate leads to slow convergence, while a large one may lead to invalid *B*^(*k*+1)^. Therefore, to select a suitable learning rate, we find the maximum λ^(*k*)^ that produces *B*^(*k*+1)^ meeting the constraint in (12) (Nocedal and Wright, 1999). This is done by applying the backtracking line search (Armijo, 1966).

To avoid local minima and to accelerate the convergence, the Gauss-Newton method is integrated with the coarse-to-fine approach (see Fig. 3). This approach first represents images with multiple resolution levels. The image representation at a coarser level is obtained simply by averaging over adjacent cells. Next, the displacement field in the coarsest level is estimated by minimizing the objective function in (12) using the image representation at this level. The estimated displacement field at the coarser level is interpolated. The interpolated result is considered as the initial guess for the optimizer at a finer level. The process of interpolation and estimation is repeated until the displacement field at the finest level is obtained. Finally, the corrected images are obtained by unwarping the distorted images with the estimated field *B*, as shown in Eq. (4). This coarse-to-fine optimization approach is summarized in Algorithm 1.

**Figure 3:**
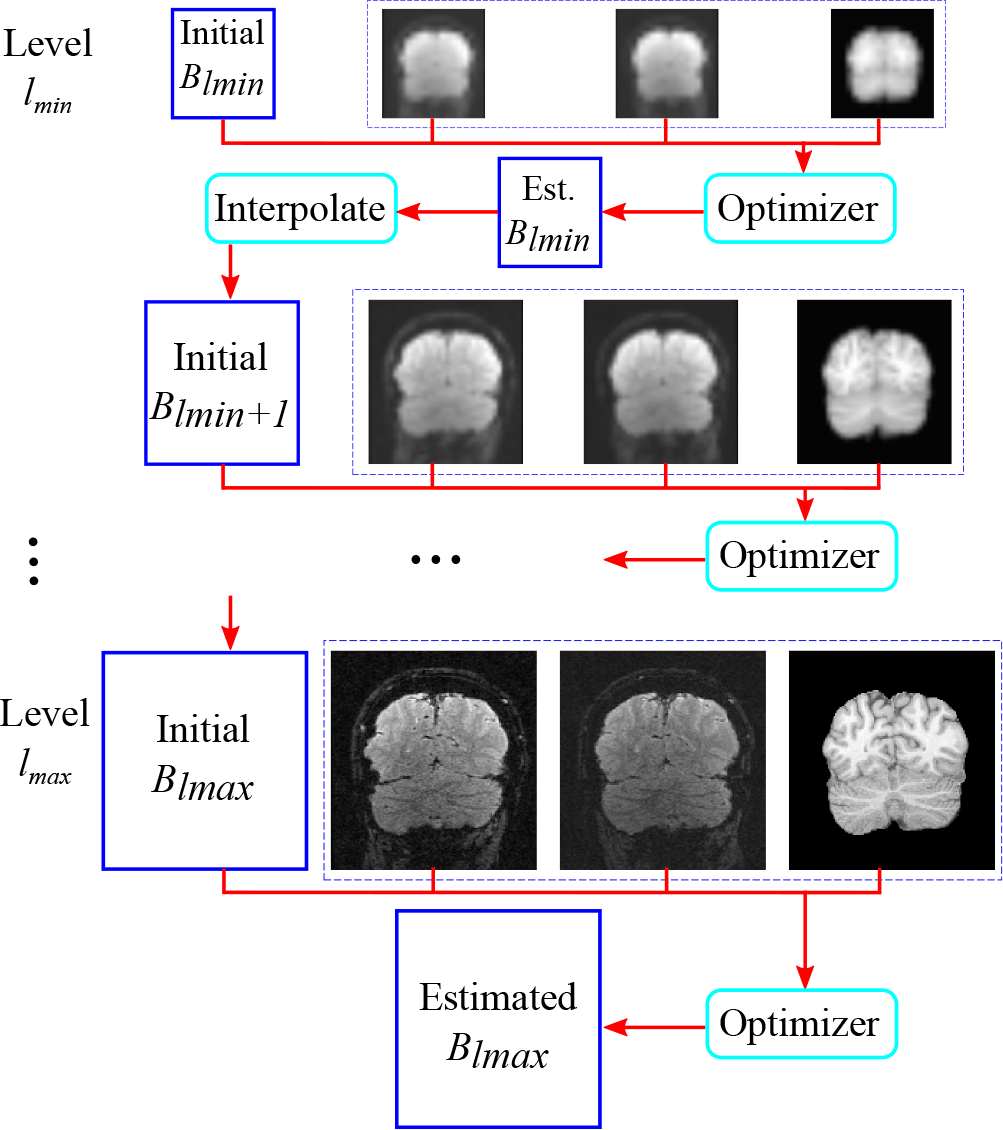
The block diagram of the coarse-to-fine optimization scheme. The displacement field is estimated at each level of data representation.

#### Algorithm 1 Coarse-to-fine Gauss-Newton for SAC

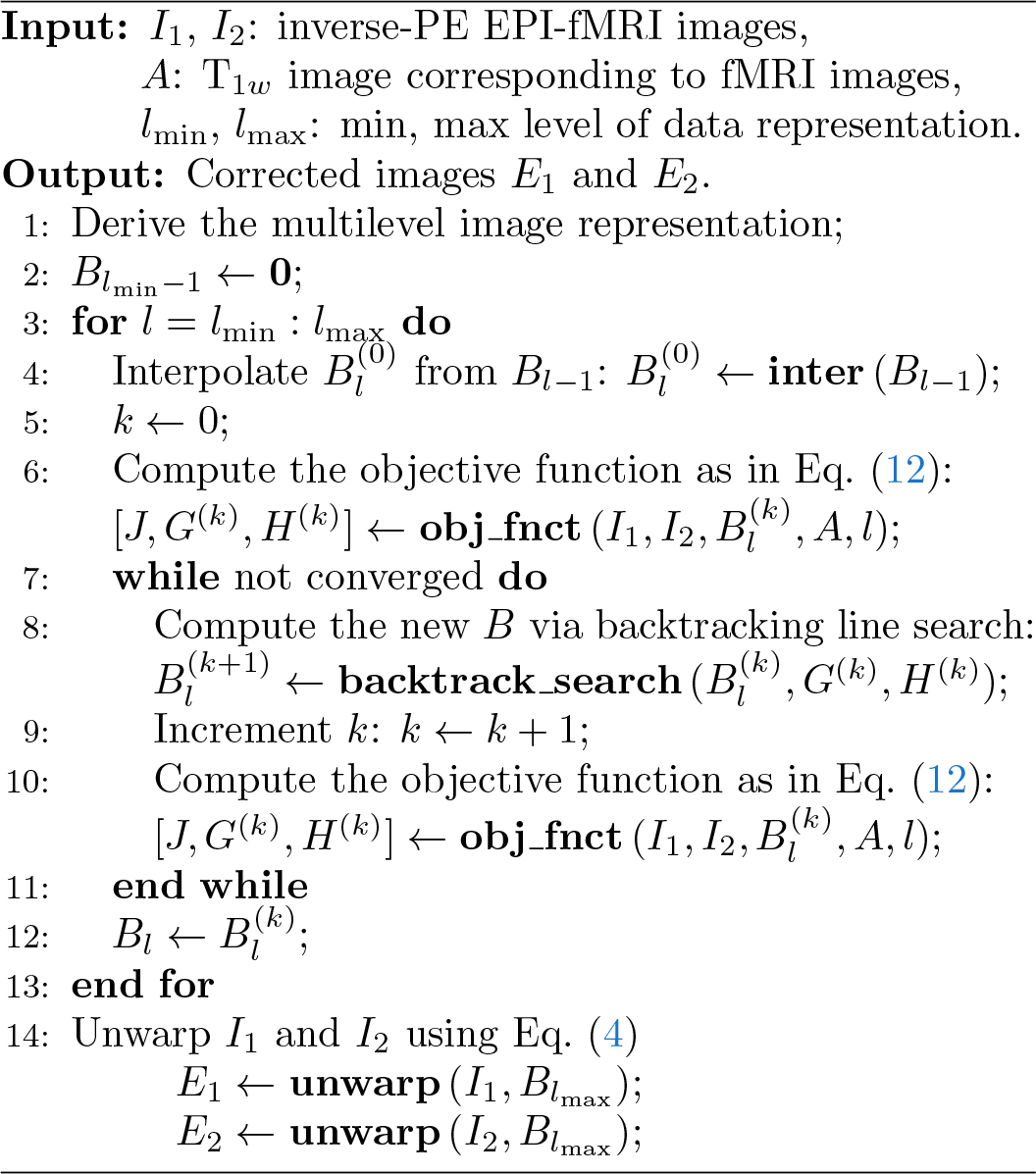

### 3.2. Optimization of hyper-parameters

The inverse-PE based SAC can be considered as an ill-conditioned inverse problem in which the choice of the regularization parameters (hyper-parameters) is crucial. Here, we propose a method to select the most suitable regularization parameters for the SAC problem. The proposed hyper-parameters optimization method is based on the Bayesian optimization (BO) with a Gaussian process (GP) prior.

The hyper-parameter optimization is performed by minimizing an error function *f*(**x**) of the given SAC method over a dataset D, where **x** is a vector of hyper-parameters. The error function here is defined by the sum of the dissimilarity measure *M* between the T_1*w*_ image and corrected fMRI images, which are produced by the SAC method given input images from the dataset D. In this study the MIND-based measure is used (refer Appendix A for a detailed description).

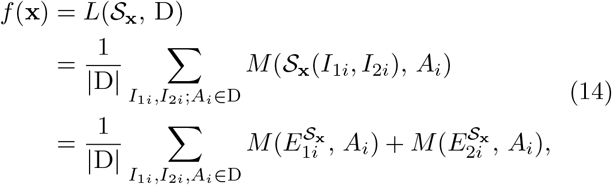

where 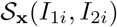 provides corrected images 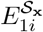 and 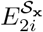 corresponding inputs *I*_1*i*_ and *I*_2*i*_, by applying the SAC method 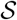 with hyper-parameters **x**. We select the hyper-parameters which give minimum error function. In other words, finding the hyper-parameters is to minimize the error function Eq. (14). Since the error function of hyper-parameters is computationally expensive, and its distribution is unknown, the hyper-parameter optimization problem becomes more challenging.

BO is a powerful technique for finding extrema of an objective function that has no closed-form expression or is computationally intensive to evaluate (Brochu et al., 2010; Bergstra et al., 2011; Snoek et al., 2012). The BO algorithm uses previous observations, which are pairs of {**x**, *f*(**x**)}, to determine what is the next optimal point for sampling the error function.

To be specific, the BO algorithm first computes the posterior expectation of what the function *f* looks like based on its previous observations. This step is done by first considering that the distribution of *f*(**x**) is a normal likelihood with noise. The error function *f* then can be considered as a GP, which is specified by the mean ***μ*** and covariance ****σ**** of a normal distribution over possible values of *f*(**x**). The means and covariances allow us to update our beliefs of what the function *f* looks like. They can be obtained by fitting the GP to a given set of observations 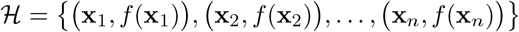.

Next, a new point is selected to sample the function *f* such that it provides a higher value of *f* or is in the unexplored region. As shown in Bergstra et al. (2011), the point can be found by maximizing the expected improvement function, which is defined as

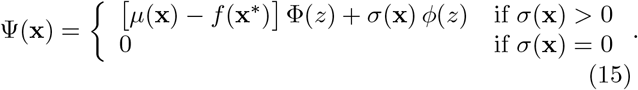

where **x**\* is the current optimal hyper-parameter point, *μ*(**x**) and *σ*(**x**) are the estimated mean and variance of function *f* at **x** in the previous step, 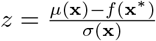, Φ(*z*) is the cumulative distribution, and *ϕ*(*z*) is probability density function of the standard normal distribution.

#### Algorithm 2 Hyper-parameters optimization algorithm.

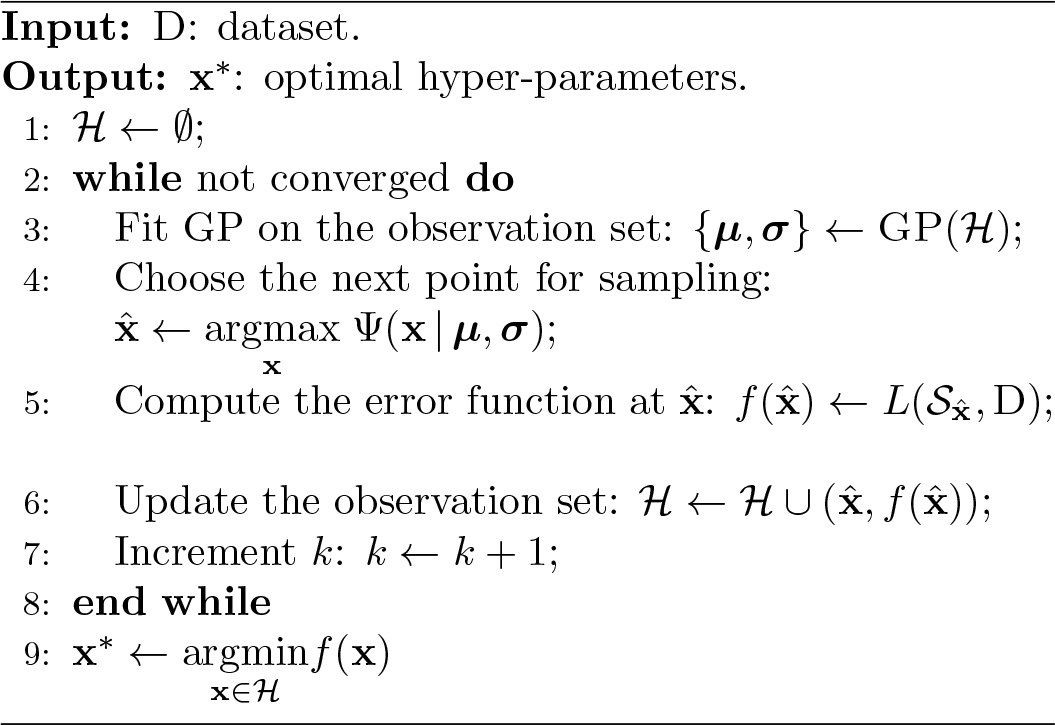

The new point obtained by maximizing the expected improvement function Ψ(**x**) is admitted to the observation set. The procedure of fitting GP and finding the sampling point is repeated until the convergence criterion is met. Algorithm 2 shows details of the Bayesian optimization for automatically selecting the hyper-parameters.

## 4. Experiments and results

In this section, 4.1 describes data acquisition and preprocessing. 4.2 presents the evaluation measures. 4.3 shows the experimental methods. 4.4 presents an analysis of the proposed method. 4.5 shows the comparison of SAC methods. 4.6 discusses the experimental results.

### 4.1. Data acquisition and preprocessing

Two EPI-fMRI datasets of the occipital cortex were used to evaluate the performance of SAC methods. The first dataset had three subjects and was acquired using a 3T scanner with an isotropic resolution of 1 × 1 × 1 mm^3^. The second dataset had three subjects and was acquired using a 7T scanner with a resolution of 0.833 × 0.833 × 0.810 mm^3^.

The 3T dataset was acquired using a Siemens 3T MAGNETOM PRISMA with a 64-channel head coil and a 2D single-shot gradient-echo (GRE) EPI sequence. Ascending and interleaved coronal slices were acquired with a repetition time (TR) of 3000 milliseconds (ms), which is also the volume repetition time, TE of 30 ms, a flip angle of 90degrees, and an image size of 192 × 144 × 36. The field of view (FOV) was 144 mm × 192 mm.

The 7T dataset was acquired using a Siemens 7T MAGNETOM whole-body research scanner with a 32-channel head coil (Nova Medical, Wilmington, US) and a 3D EPI sequence (WIP1080) (Poser et al., 2010). The sequence used a blipped CAIPIRINHA (Breuer et al., 2006; Setsompop et al., 2011), implementation (Poser et al., 2013) with the following parameters: TE of 30 ms, TR of 83ms, volume repetition of 1992 ms, flip angle of 17 degrees, echo spacing of 1 ms, FOV of 160 mm × 160 mm, matrix size of 192 × 192 × 48. The image acquisition was accelerated by a factor of 2 in-plane and by a factor of 2 in the slice-encoding direction with a CAIPI-shift of 1. This results in a total acceleration factor of 4. The image reconstruction was done by using the GRAPPA pipeline (Griswold et al., 2002), as provided by the vendor.

Figure 4 shows the three different orientation views (coronal, sagittal, and axial) of 7T inverse-PE EPI images (pink) overlaying on the T_1*w*_ image (green). The figure demonstrates that the misalignment of EPI to the T_1*w*_ image occurs mainly in one spatial direction (left-to-right).

**Figure 4:**
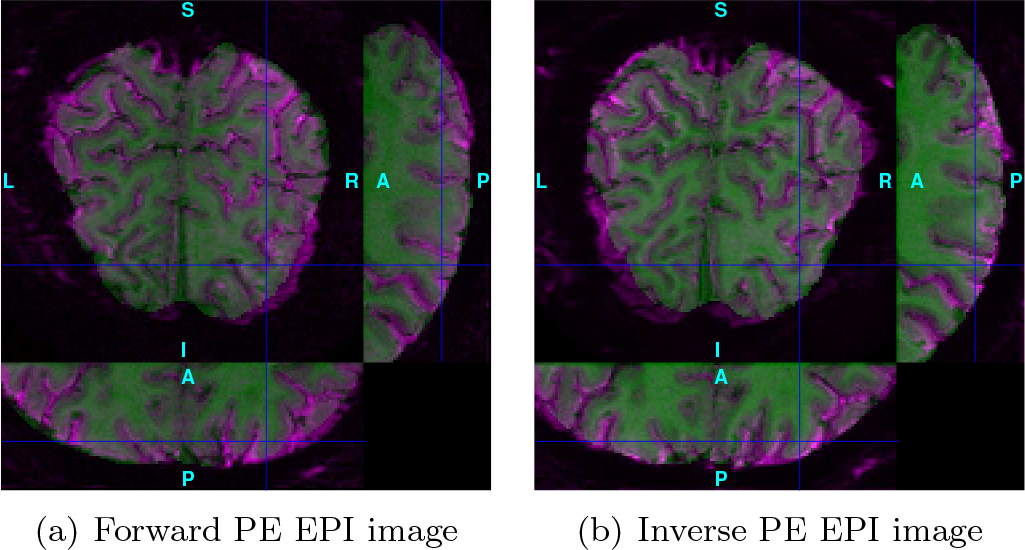
Distorted EPI slices overlaying on a T_1*w*_ image with three different orientation views. The blue lines (cross-hairs) indicate the intersection point of the three views. See the electronic color image.

Functional MRI data were acquired while subjects were presented with retinotopic mapping stimuli. In the 3T dataset, stimuli consisted of drifting bars, expanding rings, rotating bowties, and flashing full-field (see Fig. 5). Each subject took part in two scanning sessions; in each session, subjects viewed visual stimuli while scanning using either left-to-right (LR) or right-to-left (RL) blips, such that each blip accounted for half the scans. This resulted in pairs of scans with reversed patterns of distortions in the PE direction. In the 7T dataset, only the rotating bowtie stimulus was used. In each subject, two scans (with 183 or 187 volumes each) were collected with LR blip, and two short 20 s measurements with ten repeated EPI volumes were collected with the inverse blip, one at the beginning of the experimental runs and one at the end.

**Figure 5:**
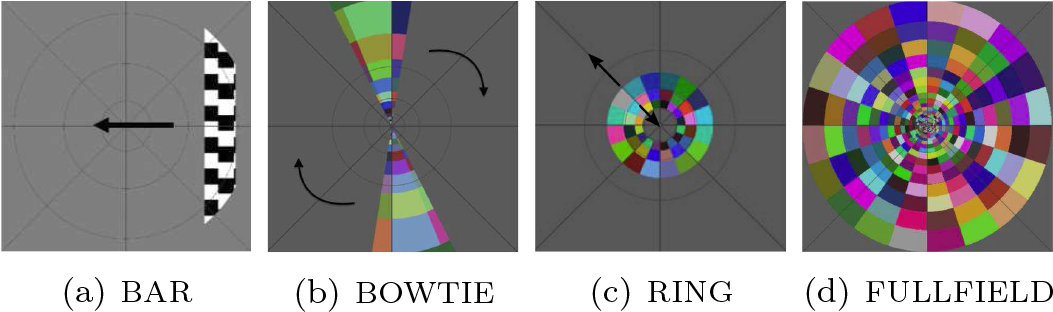
Examples of visual stimuli presented to the subjects during scanning.

For each subject in the 3T dataset, a T_1*w*_ image of the whole-brain was acquired using the 3D GRE-MRI sequence, with cubic voxels of 0.75 mm edge length. The T_1*w*_ image was then upsampled into an image with an isotropic resolution of 0.5 × 0.5 × 0.5 mm^3^. For each subject in the 7T dataset, a whole-brain anatomical image was collected using an MP2RAGE sequence (WIP900b17a) (Marques et al., 2010), with a resolution of 0.500 × 0.533 × 0.533 mm^3^.

In the first preprocessing step, all fMRI images were motion-corrected using tools in SPM12 (Penny et al., 2006). The 3T dataset was also slice scan time corrected. Hereinafter, the completely un-preprocessed (*i.e.* without motion and slice scan time corrections) images are referred to as *raw* data; and the preprocessed images are referred to as *uncorrected* or *motion correction* (MoCo) data. A T_1*w*_ alignment image of each subject was created by aligning the T_1*w*_ image to an average of two oppositely-distorted images of the subject, through SPM’s co-registration procedure (Collignon et al., 1995).

### 4.2. Performance measures

We quantitatively evaluate the corrected images in three aspects: geometric correction, blurriness, and the suitability for BOLD analysis. The various performance measures are described in this subsection.

**Anatomical similarity measures** are used to evaluate how well the corrected fMRI image matches the brain anatomy given by the T_1*w*_ image. Here, we used the mutual information (MI) to compute the similarity between the fMRI images and the T_1*w*_ image (Wells et al., 1996). A smaller value of MI indicates less similarity between the functional and anatomical images.

**The percentage of activated voxel** evaluates both the geometric accuracy and the suitability for subsequent BOLD analysis. The reason is that the BOLD response is localized in gray matter and to a certain degree in the cerebrospinal fluid (CSF) more for 3T and less for 7T data, but not in white matter. Distortions of fMRI images result in some significantly modulated voxels being mislocated in white matter of the T_1*w*_ image. Here, we employ correlation analysis, a common and robust method for analyzing phase-encoded retinotopic mapping data. This analysis provides a phase-map^5^ of the BOLD responses (Engel et al., 1997; Schira et al., 2009). Fig. 6 shows an example of a phase-map obtained by correlation analysis of an uncorrected fMRI scan. Voxels with supra-threshold response are marked in color, where the color depicts the phase (delay) of the response, not the strength of the activation. In the given example, there are many activated voxels located in white matter, indicating that they are displaced by distortions.

**Figure 6:**
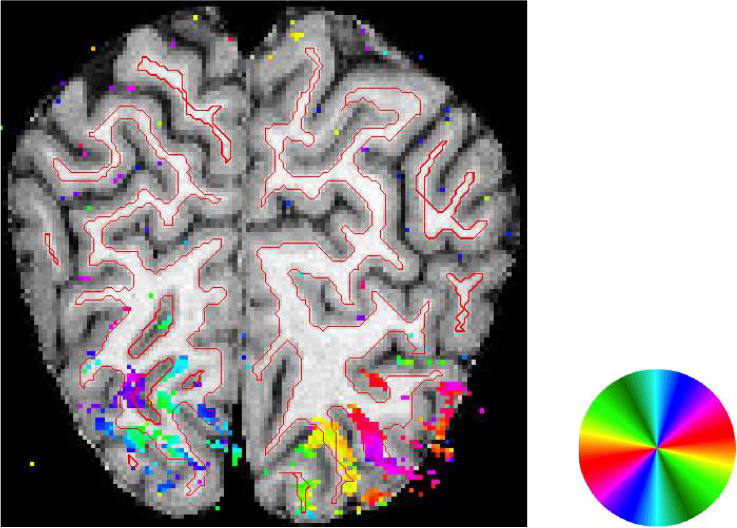
An example of a phase-map in the coronal plane. The red line marks the outer boundary of white matter. Note that the color coding represents the position in the visual field (see the color wheel), not the strength of responses, as typical in phase-encoded retinotopic mapping.

In this paper, we introduce a measure using the percentage of activated voxels in gray matter and white matter to evaluate the geometric correction in the corrected images. The reasons of measuring the percentage of activated voxels in white matter are: (i) white matter is surrounded by gray matter; (ii) there is a large number of activated voxels aligned to white matter; and (iii) it is easy to obtain an accurate and reliable segmentation of white matter from the T_1*w*_ image. The percentage of activated voxels in CSF is not considered as it is not diagnostic for geometric accuracy. A higher percentage of activated voxels in gray matter and a lower percentage of activated voxels in white matter indicates a better alignment of the fMRI images to the T_1*w*_ image.

**Blurriness measure** is used to evaluate how blurry the image is. Introducing blur to high spatial resolution fMRI data is typically undesirable (Polimeni et al., 2018; Huber et al., 2018), as it negates the often considerable effort to achieve high spatial resolution. To measure blurriness, we extended the measure proposed by Crete et al. (2007) for 2D images, to work for 3D images. This measure reflects the intensity variation of an image with respect to that of the low-pass filtered image.

The normalized intensity variation of image *I* in the *i*^*th*^ direction is defined as

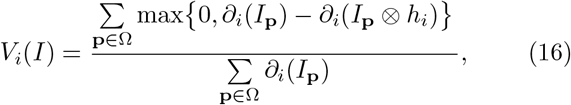

where *h*_*i*_ is a low-pass filter, and *∂*_*i*_(*I*_**p**_) is the partial derivative at point **p** in the *i*^*th*^ direction. The blurriness measure for a 3D image *I* is the sum of the normalized intensity in three directions

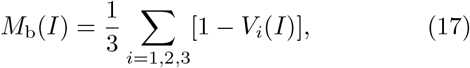

An image with a higher value of *M*_b_ is more blurred than the one with a lower value. Fig. 7 shows an example of the blurriness measure.

**Figure 7:**
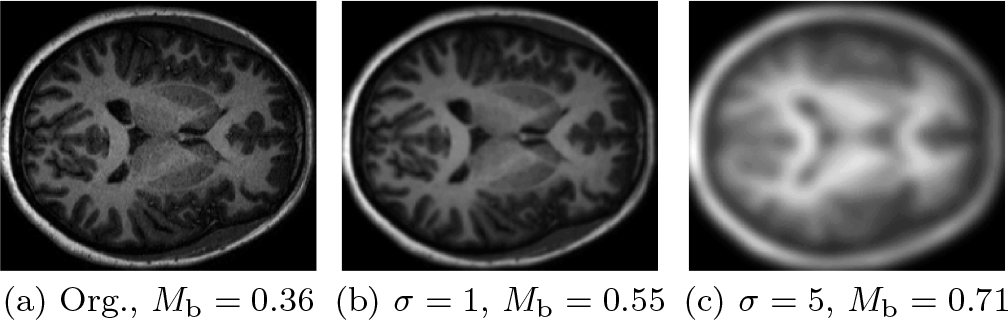
Blurriness measurements of a 3D T_1*w*_ image and blurred images by Gaussian smoothing filters with different values of standard deviation *σ*.

**Suitability for BOLD analysis** measures undesired changes in the BOLD responses. For this, we estimate the cumulative distribution function (CDF) of the phase-map values in every slice. The suitability for BOLD analysis is defined as the difference between the CDFs of corrected and uncorrected data. It is measured by the normalized cross-correlation (NCC) function. The range of NCC is in [0, 1]. A small value of NCC indicates a significant change of the BOLD responses between the corrected fMRI images and uncorrected images and vice versa.

### 4.3. Experimental methods

Scans of inverse blips were first paired together. A mean image over time of each scan was then generated. The mean images of each scan pair were processed by the SAC methods to estimate the displacement field. The estimated displacement field was then used to unwarp all volumes in the scan pair. AISAC and HySCO use the same framework implemented in MATLAB to unwarp the distorted images, while TOPUP uses another framework implemented in C. However, these unwarping frameworks are all based on the cubic spline interpolation.

We evaluated the sub-components of AISAC, which are anatomy-based registration (AR), and Bayesian optimization (BO). The tested configurations include AR and AR with BO (the complete AISAC). For settings without BO, the regularization parameters were tested then set as *α* = 30, *β* = 50, and *γ* = 75000.

We further compared the proposed AISAC method with two state-of-the-art SAC methods: HySCO (from the SPM12 toolbox version r7219) and TOPUP (from the FSL package version 5.0.9). For each pair of inverse-PE scans, the displacement field was estimated using two mean images of these scans and then used to unwarp the distorted images from these scans. The regularization parameters of AISAC were selected automatically by applying the BO technique, while these parameters in HySCO were set as *α* = 50 and *β* = 10 as suggested in Ruthotto et al. (2012). The regularization parameters of TOPUP were selected as indicated in the preprocessing pipeline for the HCP (Glasser et al., 2013).

We also assessed the time complexity of three SAC methods by recording their execution time with inputs as pairs of mean images. All timing reported were collected on a Linux workstation with an Intel Xeon Processor E3-128V2 3.6 GHz and 32 GB RAM.

### 4.4. Analysis of the proposed method

We investigated whether the BO technique improves the anatomy-based registration scheme. Table 2 shows the similarity measures and execution time of the two AISAC settings. It appears that all tested settings provided corrected images with comparable quality, *i.e.* similarity to the anatomical image. However use of the BO technique resulted in a faster run time than when it was not used.

**Table 2:**
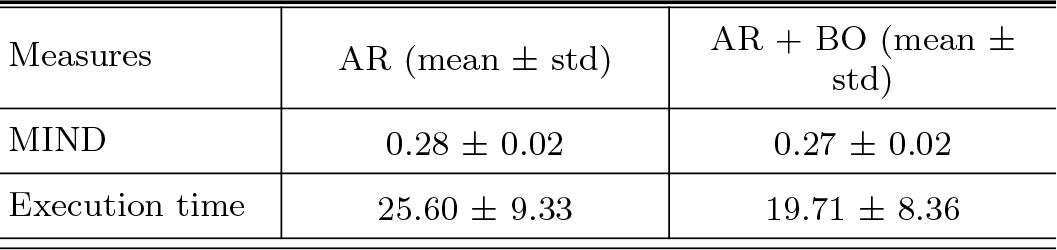
Similarity measures and execution time (in seconds) of AISAC configurations: with and without using the BO technique. Methods using BO do not include time for estimating the hyper-parameters.

### 4.5. Comparisons of SAC methods

First, we investigated the time complexity of the proposed method. Table 3 shows the mean computation time of three SAC methods: AISAC, HySCO, and TOPUP. Overall, AISAC is the least computationally expensive, about 1.4 times faster than HySCO and over 20 times faster than TOPUP.

**Table 3:** Execution time of SAC methods (in second).

| Datasets | TOPUP (mean ± std) | HySCO (mean ± std) | AISAC (mean ± std) |
| --- | --- | --- | --- |
| 3T dataset | $399.87 \pm 6.52$ | $21.19 \pm 5.38$ | $14.88 \pm 1.78$ |
| 7T dataset | $741.33 \pm 6.04$ | $43.88 \pm 12.03$ | $32.58 \pm 3.80$ |

Second, we visually assessed the quality of corrected images generated by the same three SAC methods. Fig. 8 shows the uncorrected and the corresponding corrected images of these SAC methods tested from two subjects, one in the 3T dataset (top row) and the other in the 7T dataset (bottom row). The corresponding T_1*w*_ images are presented in the right-most column. All tested SAC methods decreased SA distortions noticeably. In both datasets, the AISAC method produced sharp images with clearly visible tissue interfaces, especially near the brain-air interface, while TOPUP produced low contrast images and HySCO produced images with artifacts in the brain-air interface (see blue arrows).

**Figure 8:**
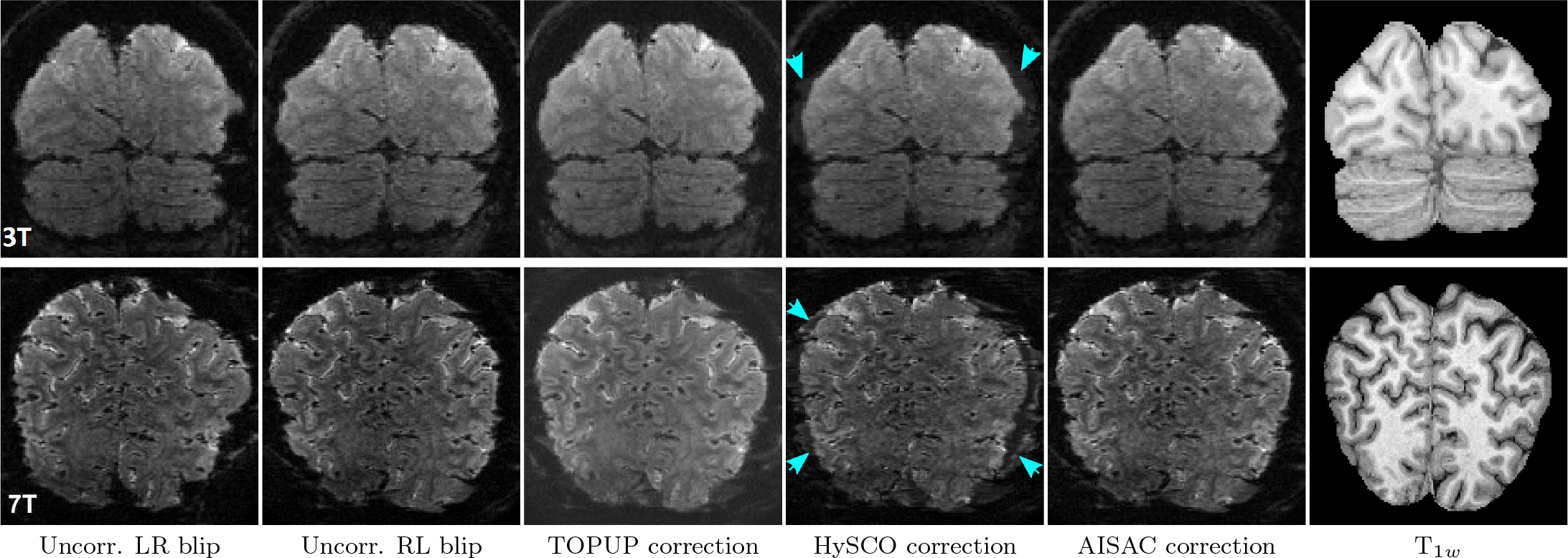
Uncorrected images and their corrected versions created using three SAC methods and corresponding T_1*w*_ images. The top row presents images of a subject in the 3T dataset. The bottom row shows images of a subject in the 7T dataset. Arrows points the artifacts produced by HySCO.

Third, we analyzed the level of blurriness that each SAC method produces. The left column in Fig. 9 shows the cumulative distribution of the blurriness measure over images of each dataset, comparing uncorrected (MoCo) data with corrected data (SACs applied after MoCo). In general, all SAC methods introduced blur when compared to uncorrected images. HySCO produces the most blur in both datasets. AISAC produces the least blur, especially in the 3T dataset. Next, we aimed to assess the amount of blur introduced by AISAC correction by comparing the blurriness measure for: (i) MoCo, (ii) AISAC after MoCo, and (iii) blurred (Gaussian filtered with *σ* = 0.3) MoCo images. The middle column in Fig. 9 shows the cumulative distribution of these measures. These graphs indicate that AISAC blurs the images with the same degree as applying a Gaussian filter of *σ* = 0.3 (FWHM = 0.7) of the voxel size. We also compared the amount of blurriness that motion correction and AISAC correction introduce to the raw data. The right column in Fig. 9 shows cumulative distributions of the blurriness measure from five image types: (i) MoCo, (ii) AISAC after MoCo, (iii) raw images, and (iv) and (v) blurred version of raw images by Gaussian filters. These graphs show that motion correction blurs the raw images with the same degree as applying a Gaussian filter of *σ* = 0.37 for 3T and of *σ* = 0.33 for 7T (see purple dotted lines). Motion correction and SAC blurs the raw images with the same degree as applying a Gaussian filter of *σ* = 0.38 for 3T and of *σ* = 0.35 for 7T (see green dash-dot lines). This suggests that AISAC introduces a much smaller degree of blurriness than the motion correction procedure or any other SAC correction method.

**Figure 9:**
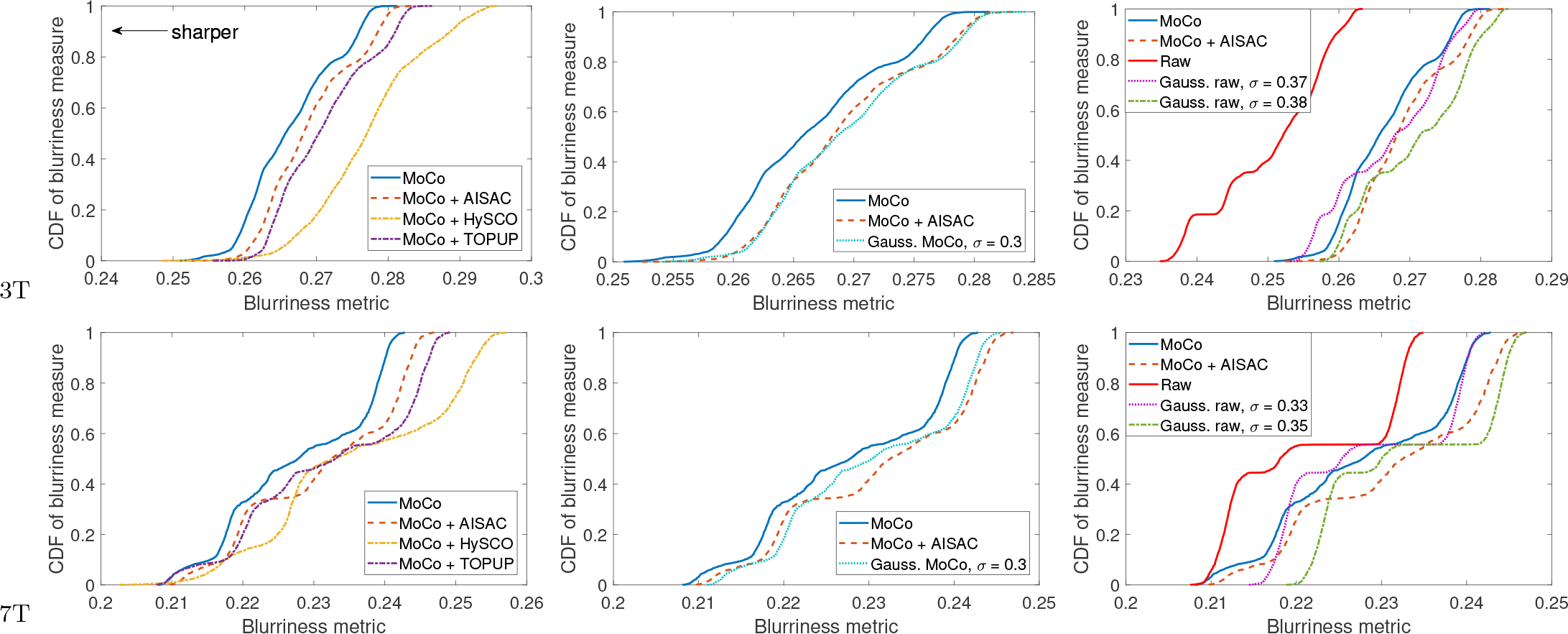
Cumulative distributions (CDs) of the blurriness measures over subjects for the 3T (top row) and the 7T (bottom row) datasets. The left column shows CD graphs of uncorrected (MoCo) and corrected images after MoCo. The middle column shows CD graphs of MoCo images, AISAC corrected images and blurred version of MoCo by a Gaussian filter with *σ* = 0.3. The right column shows CD graphs of MoCo images, AISAC corrected images, raw images, and blurred versions of raw images by Gaussian filters.

Fourth, we computed the anatomical similarity measures for more quantitative evaluation. Fig. 10 shows box-plots of MI coefficient over images of the 3T dataset (6140 images) and the 7T dataset (1667 images). The MI results show that the corrected images are more similar to the anatomical image than the uncorrected images. However, these MI coefficients do not show clear differences among SAC methods, especially for the 7T dataset, failing to capture the differences which are visible to the human observer.

**Figure 10:**
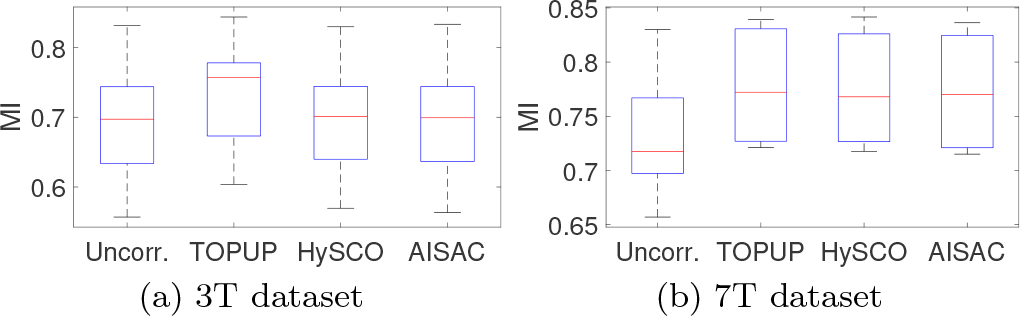
Box-plots of the MI coefficients between anatomical and fMRI images.

Fifth, we tested if AISAC improves the accuracy of geometric distortion in fMRI analysis. Fig. 11 shows phase-maps of uncorrected and three SAC corrected data of a subject in the 7T dataset, with the coronal and axial views. More phase-map examples comparing SAC methods are shown in Figs. B.14 through B.15. Visual inspection reveals that distortions are smaller in the 3T datasets than the 7T datasets, where uncorrected 7T data exhibits clear misalignment between activated voxels and gray matter. The maximum misalignment is 5 pixels (about 4.16 mm, see arrows on the phase-maps of uncorrected data in Fig. 11). Also, visual inspection suggests that AISAC correction results in better alignment than the competing methods (though only marginally better than TOPUP).

**Figure 11:**
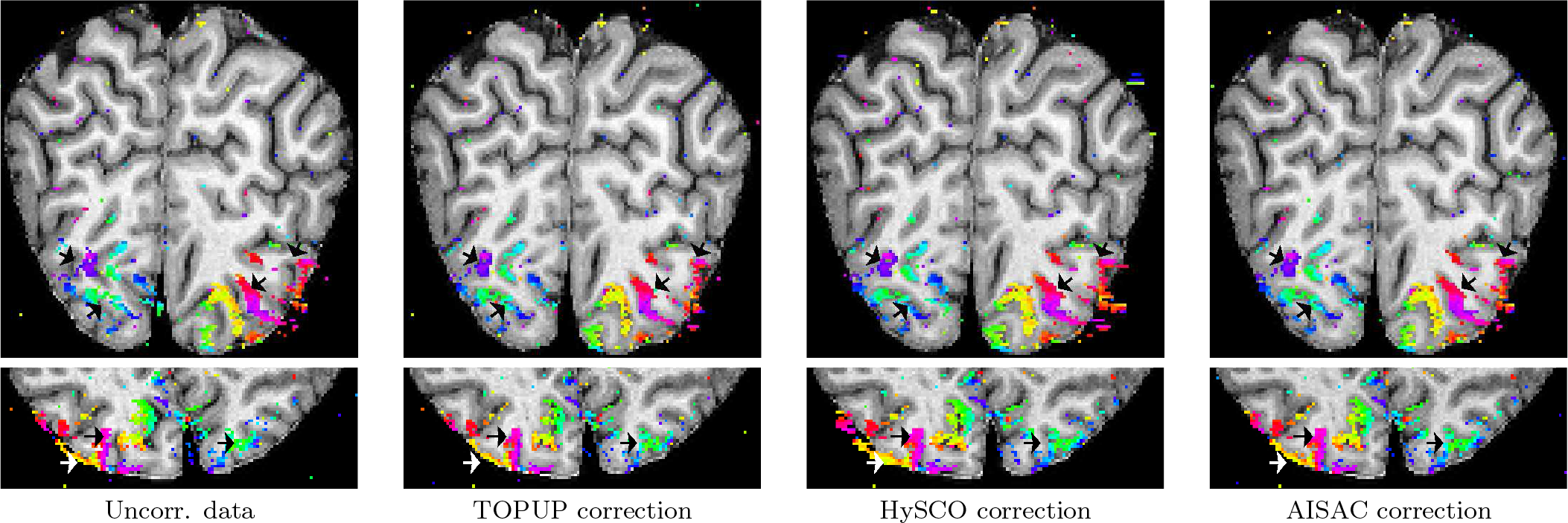
Phase-maps projected onto the T_1*w*_ image of uncorrected and corrected data in the 7T data. The top row shows phase-maps in the coronal view. The bottom row shows phase-maps in the axial view. Arrows point to the areas with large distortions. See the electronic color image.

To quantify this, we calculated the percentage of activated voxel measures. Fig. 12 shows the percentage of activated voxels (PAV) in gray matter and white matter in 3T and 7T datasets. Table 4 shows the mean number of activated voxels, the number in gray matter and white matter, and the percentage of activated voxels. In the 3T dataset, the SAC methods do not improve the PAV in gray matter but decrease the PAV in white matter. In the 7T dataset, the SAC methods both improve the PAV in gray matter and decrease the PAV in white matter. In general, all three SAC methods greatly improve the alignment of BOLD response to gray matter, especially in datasets with large geometric distortions, and AISAC improves the alignment the most, with the highest PAV in gray matter and lowest PAV in white matter.

**Table 4:** Number of activated voxels of data over scans.

| Datasets | Activated voxel types | Uncorr. | TOPUP | HySCO | AISAC |
| --- | --- | --- | --- | --- | --- |
| 3T | All | 7475 | 7628 | 7861 | 7747 |
|  | In GM | 2126 | 2066 | 2057 | 2066 |
|  | In WM | 304 | 304 | 310 | 288 |
| 7T | All | 6143 | 6681 | 7014 | 6737 |
|  | In GM | 3236 | 3910 | 4016 | 4093 |
|  | In WM | 1392 | 725 | 922 | 572 |

**Figure 12:**
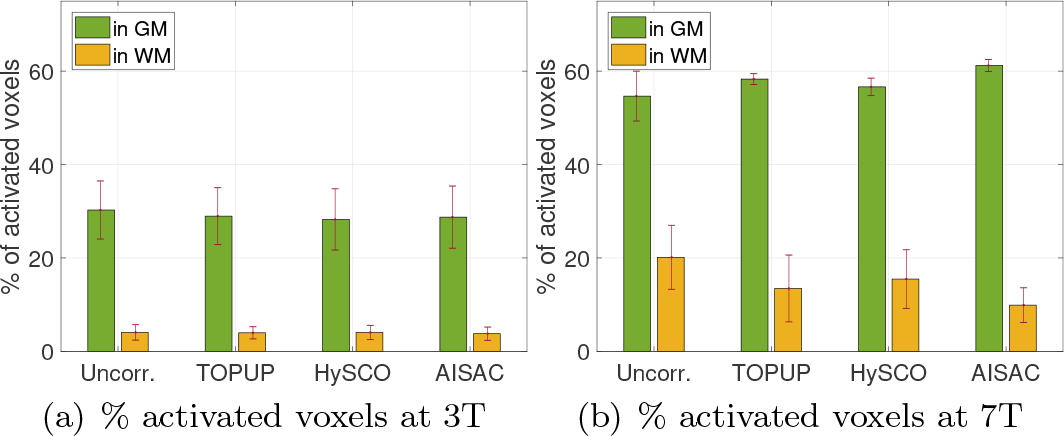
Mean percentage of activated voxels in gray matter and white matter. The error bar shows the corresponding standard deviation of the percentage.

Finally, we evaluated the suitability for BOLD analysis. Table 5 shows the mean and standard deviation of the NCC between estimated CDFs of the phase values before and after applying SACs. SAC methods should maintain the BOLD responses. The results in Table 5 indicate that the BOLD responses of all three correction methods are not different from the one of uncorrected data.

**Table 5:** Change of the BOLD responses after SACs over scans.

| Datasets | TOPUP<br>(mean ± std) | HySCO<br>(mean ± std) | AISAC<br>(mean ± std) |
| --- | --- | --- | --- |
| 3T | $0.989 \pm 0.042$ | $0.970 \pm 0.073$ | $0.970 \pm 0.073$ |
| 7T | $0.998 \pm 0.009$ | $0.999 \pm 0.004$ | $0.999 \pm 0.004$ |

### 4.6. Discussion

The experimental results indicate that SAC methods can correct geometric distortions in EPIs, even when these distortions are severe, such as in the 7T dataset. HySCO produces corrected images with ghost artifacts around the brain boundary, while TOPUP produces images with good distortion corrections and no obvious artifacts but introduces blur and affects the BOLD responses. Judged by visual inspection and the performance measures, AISAC correction provides the images with the best alignment to the anatomical image, compared to TOPUP and HySCO, especially in the dataset with severe geometric distortions.

For high spatial resolution fMRI, the blurring effects on post-processing are of great concern. We found that all the SAC methods introduce blur into corrected images, but the proposed AISAC method introduces the least amount. Furthermore, the blur that SAC methods introduce to the motion-corrected data is much less than the blur that the motion correction introduces to the raw data.

The anatomical similarity results indicate that the mutual information measure is able to reflect obvious improvements between uncorrected and corrected images. However, the MI measure is not suitable to evaluate the anatomical similarity of corrected images, whose differences are not apparent. A possible reason is that MI uses the distribution of the intensity, and it is not sufficient to capture the small differences between images; his leads to a mismatch between the MI measure and human visual inspection.

## 5. Conclusion

This paper introduced a novel method, AISAC, for correcting susceptibility artifacts in high spatial resolution EPI-fMRI images. The proposed method uses inverse-PE EPI-fMRI images and the T_1*w*_ image to register the distorted EPI-fMRI image. The symmetric registration principle is used in the proposed AISAC method, where the corrected image is considered as a middle of inverse-PE images and as similar to the T_1*w*_ image as possible. The utilization of the T_1*w*_ image serves two purposes: (i) to regularize the registration for producing more robust and sharper corrected images, and (ii) to guide the selection of regularization parameters through the Bayesian optimization framework.

The performance of AISAC and two other SAC methods was evaluated using two high spatial resolution EPI-fMRI datasets. The experimental results show that AISAC outperforms the existing methods in terms of accuracy and robustness, particularly in sub-millimeter images obtained by the high field scanner. The proposed method produces sharper corrected images with better geometric correction. It is effective in preserving the structure of the T_1*w*_ structural image in regions of significant distortions. Besides, it requires less computational resources than both TOPUP and HySCO methods. The AISAC corrected data provides better results in subsequent fMRI analysis, while still keeping the BOLD responses as found in the uncorrected data. The proposed method is readily applicable to other types of fMRI data acquired by GRE-EPI.

## Acknowledgment

The authors acknowledge the facilities of the National Imaging Facility at the Center for Advanced Imaging, University of Queensland. The authors also thank Siemens Healthcare for providing the prototype WIP1080 for data acquisition. This research was supported by two grants (DP140101833 and DP170101778) from the Australian Research Council and a Matching scholarship from the University of Wollongong.

## Appendix A. Multi-modal similarity measure

Modality independent neighborhood descriptor (MIND) is a multi-dimensional descriptor, which was proposed for computing the dissimilarity measure in multi-modal deformable image registration (Heinrich et al., 2012). This descriptor is independent of the modality, contrast, and noise level of images by capturing the self-similarity of the image patches around a voxel.

The multi-dimensional descriptor *s*_MIND_ of a voxel **x** within the search space 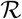 (centered at **x**) is a vector with the length as the number of elements in 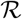. The MIND of **x** in an image *I* at a single entry 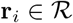 is defined as

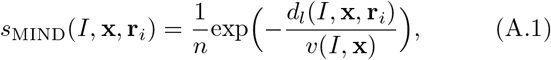

where *n* is a constant to normalize the maximum value as 1. 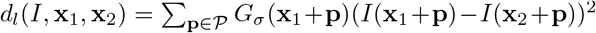 is the patch-based dissimilarity, with 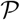 denoting neighborhood indexes of a patch size of (2*l* + 1) and *G*_*σ*_ denoting a Gaussian filtered image of the difference between image *I* and its shifted version from **x**_1_ to **x**_2_, with a kernel size as of the patch. And *v*(*I*, **x**) is the mean patch-based dissimilarity of voxel **x** in image *I* with its six neighbors.

The MIND-based dissimilarity of images *A* and *B* is defined as the sum of MIND difference at every voxel in the image domain Ω, 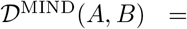 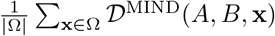 with

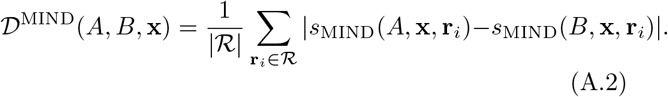

A smaller value of the MIND-based measure indicates more structural similarity between images.

Fig. A.13 shows an example of MIND difference maps (blue: low difference, red: high difference) of monomodality (T_1*w*_ images of different subjects) and multimodality (T_1*w*_ and EPI-fMRI images of the same subject) images. The first row shows slices of images to be computed MIND measure with the image shown in the first column. The second row shows the corresponding MIND difference maps. It can be seen that the T_1*w*_ images with the same intensity distribution, but different structures, have a higher MIND-based dissimilarity measure than multimodal images with the same structural information.

**Figure A.13:**
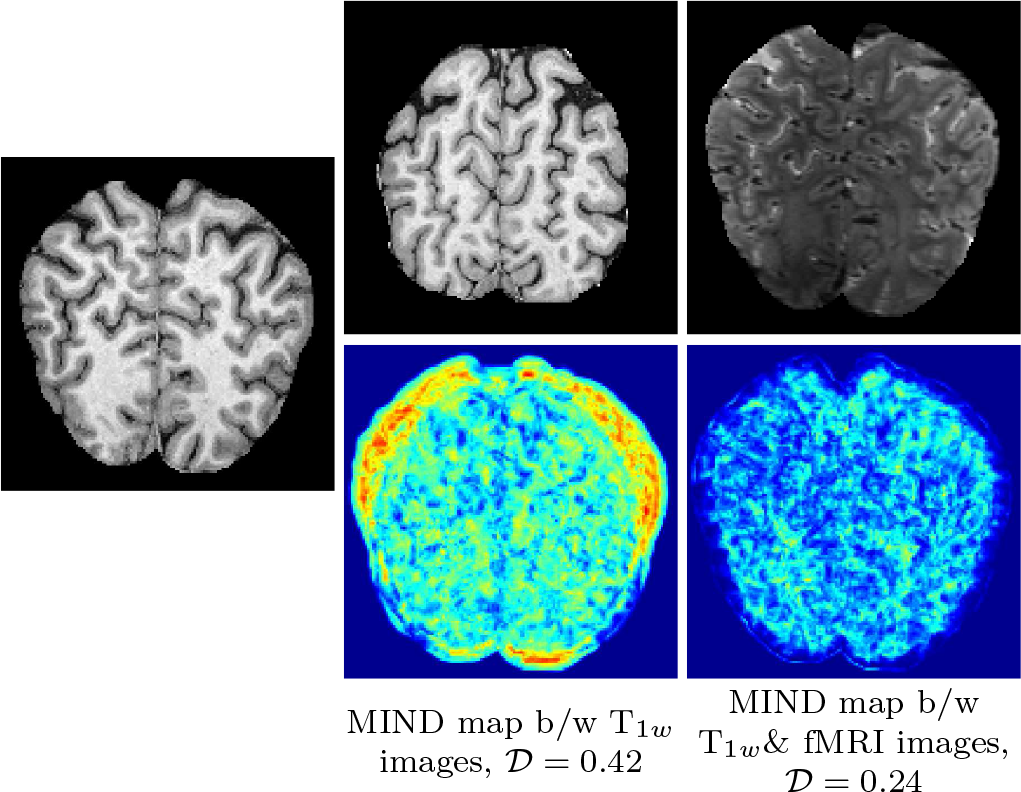
An example of MIND difference maps between T_1*w*_ (first column) and other MRI types. The first column shows a slice of a T_1*w*_ image (A). The second column shows a slice of a T_1*w*_ image from another subject (top row) and its MIND difference map with A (bottom row). The third column shows a slice of an EPI image from the same subject of A (top row) and its MIND difference map with A (bottom row). In MIND difference maps, blue color denotes a low difference, and red color denotes a high difference.

## Appendix B. Supplementary figures

**Figure B.14:**
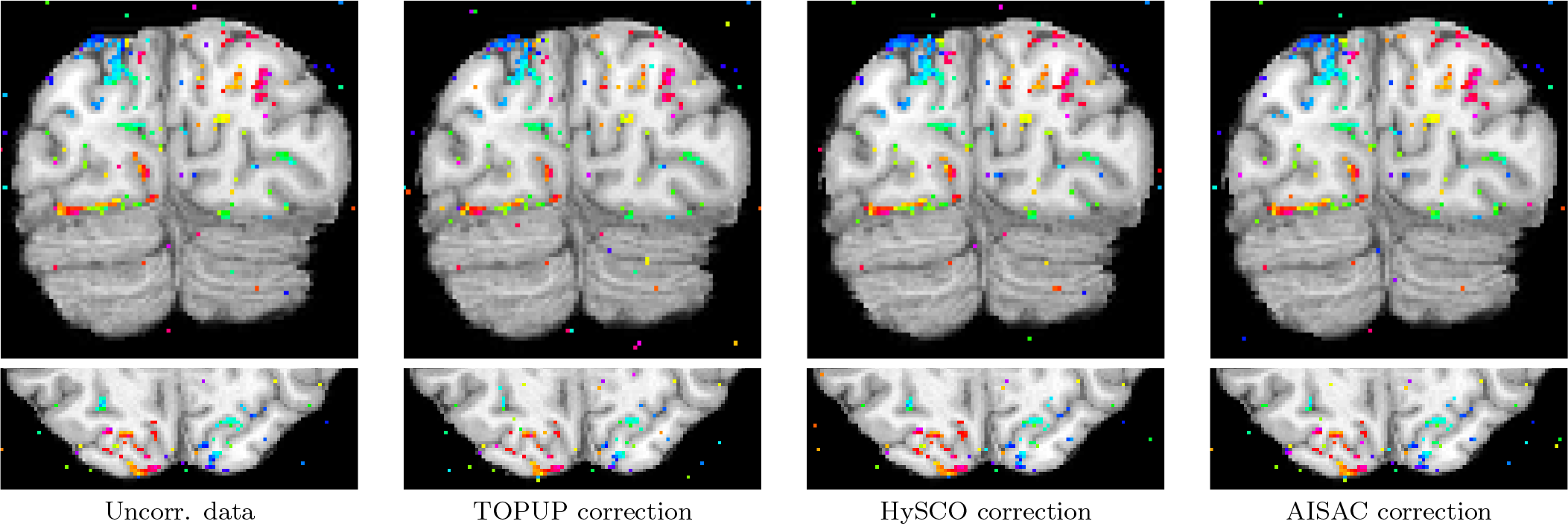
Phase-maps projected onto the T_1*w*_ image of uncorrected and corrected data in the 3T data. The top row shows phase-maps in the coronal view. The bottom row shows phase-maps in the axial view. See the electronic color image.

**Figure B.15:**
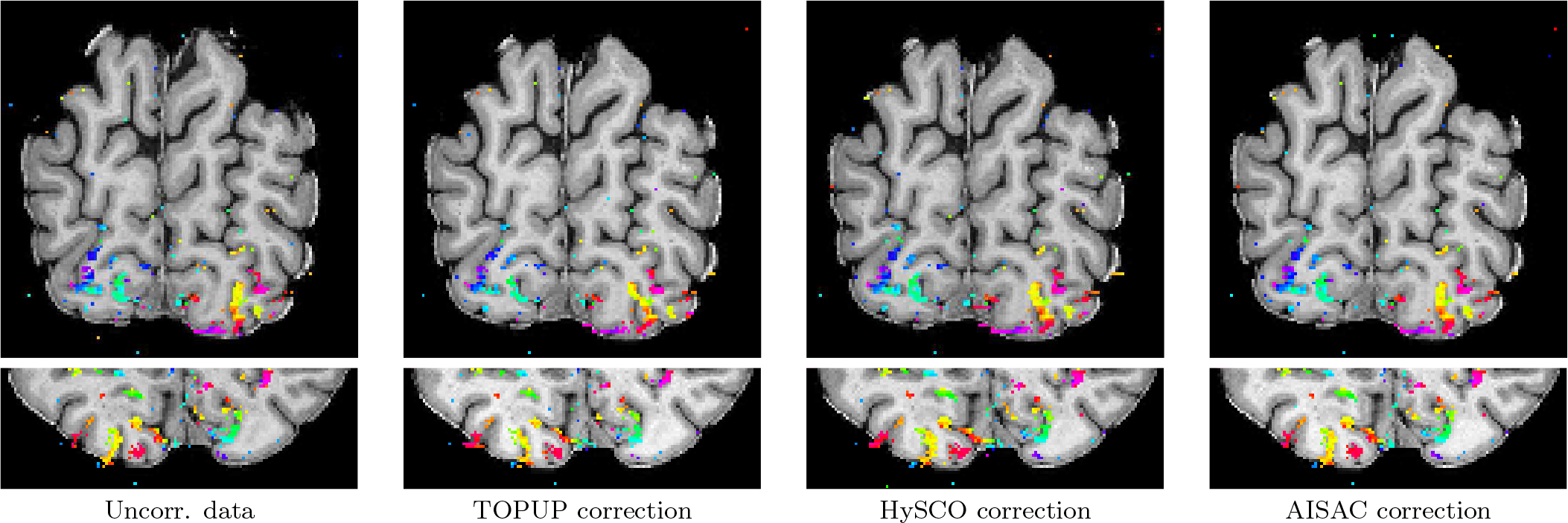
Phase-maps projected onto the T_1*w*_ image of uncorrected and corrected data in the 7T data. The top row shows phase-maps in the coronal view. The bottom row shows phase-maps in the axial view. See the electronic color image.

## Footnotes

1 In fMRI, the phase encoding direction is also known as the *polarity of phase-encoding gradient* or the *blip*.

2 https://fsl.fmrib.ox.ac.uk/fsl/fslwiki/topup

3 http://www.diffusiontools.com/documentation/hysco.html

4 The deformation grid is the sum of the regular grid and the displacement field.

5 The term phase-map refers to the use in phase-encoded retinotopic mapping, which is provided by an FFT-based analysis procedure of BOLD time courses. It is different from “phase” in the phase-encoding direction derived from k-space in MRI acquisition.

